# Conditional Myeloid-Specific Inhibition of UBE2N Hinders YUMM1.7 Growth

**DOI:** 10.64898/2026.08.31.748234

**Authors:** Kelsie Schiavone, Adam Pecoraro, Afsa Khawar, Kathleen Zhang, Daniel T. Starczynowski, Jennifer Y. Zhang

## Abstract

The role of UBE2N in myeloid cell-mediated immune suppression in cancer remains undefined. Here, we examined the function of UBE2N in myeloid cell-mediated tumor progression using a temporally inducible myeloid-specific knockout model (*LysM^CreER^Ube2n^fl/fl^*). Temporally induced deletion of *Ube2n* in myeloid cells (*Ube2n^MyeKO^*) significantly hindered growth of YUMM1.7 melanoma. This was accompanied by reduced myeloid cell burden within the tumor microenvironment. We observed altered abundance of PD-1, PD-L1, and SPP1 in the *Ube2n^MyeKO^* tumor microenvironment at the tissue level. In vitro analysis showed that knock-in expression of a catalytically deficient UBE2N^C87S^ mutant in bone marrow-derived macrophages (BMDMs) markedly decreased expression of *Spp1*. We observed decreased SPP1 secretion in *Ube2n^MyeKO^* BMDM-conditioned media (CM). Treatment with *Ube2n^MyeKO^* BMDM-CM decreased co-expression of PD-1, TIM-3, and LAG-3 on chronically stimulated T cells. Antibody-mediated neutralization of SPP1 in Ube2n^WT^ BMDM-CM decreased PD-1 expression on CD8+ T cells. Together, these findings suggest a role for myeloid UBE2N in YUMM1.7 progression.

## INTRODUCTION

Tumor-associated macrophages (TAMs) are among the most abundant immune populations in melanoma. TAMs play complex, context-dependent roles in tumor progression [1]. They exist along a functional spectrum between pro-inflammatory and immunosuppressive phenotypes, with the latter typically enriched in advanced melanomas [2]. High densities of immunosuppressive TAMs correlate with poor response to immune checkpoint inhibitor (ICI) therapy, reduced overall survival, and adverse clinicopathologic features such as mesenchymal transformation in patients with cutaneous melanoma [3,4]. Early in tumor development, macrophages may contribute to antigen presentation and immune priming, but melanoma-derived cytokines, growth factors, and hypoxia can progressively skew them toward tumor-promoting states [1,2]. In human primary and metastatic melanoma samples, accumulation of immunosuppressive macrophages increases with disease stage and is linked to enhanced invasion and metastatic dissemination [5]. Single-cell transcriptomic analyses have revealed a striking degree of macrophage heterogeneity beyond the classical M1/M2 framework, with specific transcriptional subsets independently predicting poor clinical outcomes [6]. In mouse melanoma models, experimental enrichment of immunosuppressive macrophages accelerates tumor growth, whereas depletion or reprogramming toward immunologically active states suppresses it [4,7]. Mechanistically, TAMs promote melanoma progression by driving angiogenesis, extracellular matrix remodeling, immune checkpoint expression, and suppression of cytotoxic T cell activity [2,8]. Accordingly, therapeutic strategies targeting monocyte/macrophage recruitment, expansion, or polarization are being investigated to improve responses to immunotherapy and limit melanoma progression [1,7]. However, there remains a major knowledge gap in understanding and manipulating the effectors of myeloid/macrophage-driven immune suppression.

UBE2N/UBC13 is an E2 ubiquitin-conjugating enzyme primarily responsible for catalyzing lysine-63–linked ubiquitination (K63-Ub). K63-Ub modulates protein stability and function in a non-proteolytic manner, orchestrating signal transduction, DNA repair, and cell survival [8,9]. Our previous studies have demonstrated that UBE2N promotes melanoma growth in both cancer cell-intrinsic and extrinsic manners. UBE2N directly promotes melanoma growth through MEK/FRA1/SOX10 signaling in cancer cells [10]. Other studies have demonstrated dominant roles for UBE2N in driving prostate cancer, non-small cell lung cancer, neuroblastoma, and acute myeloid leukemia (AML) [12–16]. UBE2N promotes these malignancies through proteoglycan modification, inhibition of cell apoptosis, and promotion of oncoprotein stability, as well as modulation of MAPK superfamily signaling [15]. The broad relevance of UBE2N to various cancers has led to the active development of small-molecule inhibitors. Among these, NSC697923 (NSC) inhibits the formation of UBE2N-ubiquitin thioester bonds and treatment of NSC induces malignant cell apoptosis by altering p53 and MEK signaling pathways [15]. UC764864 and its derivative UC764865 (UC65) exert steric inhibition by blocking the transfer of ubiquitin moiety from UBE2N to target proteins. When administered to mice, UC65 demonstrates a low toxicity profile and effectively inhibits progression of acute myeloid leukemia without inducing other deleterious effects [16]. Taken together, these findings establish UBE2N as a potential target for cancer therapeutics.

UBE2N also regulates immune homeostasis. Deletion of *Ube2n* in regulatory T-lymphocytes (Treg) during development impairs Treg function and sensitizes animals to early onset of autoimmune disorders [17]. Deletion of *Ube2n* in myeloid cells has little effect on macrophage development but affects macrophage phenotypic response to MyD88-dependent signaling [18,19]. Recently, we demonstrated that temporally controlled deletion of *Ube2n* in Tregs in adult mice induced a potent antitumor immunity [11]. UBE2N’s role in myeloid immune cell function has yet to be explored in the context of melanoma. In this study, we examined the role of myeloid cell UBE2N in a subcutaneous syngeneic melanoma model. We genetically targeted *Ube2n* in myeloid cells (*Ube2n*^MyeKO^) in a temporally controlled manner using the *LysM^CreER^Ube2n^fl/fl^* mouse model. We showed a significant decrease in YUMM1.7 tumor growth upon induction of *Ube2n* knockout in myeloid cells. Immunofluorescent staining analysis showed that CD11B+ myeloid cell burden significantly decreased in *Ube2n^MyeKO^* tumors, alongside decreased PD-1 co-occupancy in the CD3 compartment. Further, *in vitro* analysis showed that deletion of *Ube2n* in myeloid cells altered expression of pro- and anti-inflammatory mediators *Vegf*, *Spp1*, *Tnfa,* and *Apoe.* These markers are associated with angiogenesis [20,21], T-cell exhaustion [22,23], aggressive tumor behavior [24,25], and interference with cytotoxic T-cell activity [26,27], respectively. Finally, *in vitro* co-culture experiments suggested a role for myeloid UBE2N in regulating SPP1 secretion, with consequences for T cell checkpoint marker expression.

## MATERIALS AND METHODS

### Ethics statement

All animal experiments were done in compliance with protocols approved by the Institutional Animal Care and Use Committee. All animals were housed in a temperature and humidity-controlled facility with a 12–hour light/dark cycle with food and water continually available. Animals were assigned randomly to different treatment groups.

### Murine tumor model

*LysM^CreERT2^* mice (Strain # 032291) were purchased from the Jackson Laboratory (Bar Harbor, ME). *Rosa26^CreERT2^.Ube2n^fl/fl^* mice were kindly gifted by Dr. Shao-Cong Sun (University of Texas MD Anderson, Houston, TX) with the permission of Dr. Shizuo Akira (Osaka University, Osaka, Japan). After crossing with C57BL6 mice to remove *Rosa26^CreERT2^ alleles*, *Ube2n^fl/fl^*were crossed with *LysM^CreERT2^* mice to generate *LysM^CreERT2^*.*Ube2n^fl/fl^* mice. Tumor implantation was performed on 8–12-week-old *LysM^CreERT2^*.*Ube2n^fl/fl^*and WT or heterozygous littermates. Both male and female mice were used, and no apparent sex differences were observed. The mid-back region of each animal was shaved and injected subcutaneously (SubQ) with YUMM1.7 (1 × 10^5^) cells suspended in 100 µL sterile phosphate buffered saline (PBS). Body weight was recorded on days 0,4,7,10,13, and 16. Tumors were not yet palpable at day4 in either group, and tumor volumes were therefore recorded beginning on day 7. To induce *Ube2n* deletion, tamoxifen was dissolved in sterile filtered corn oil at a 20 mg/mL concentration and intraperitoneally (i.p) injected at a dose of 100 mg/kg of mouse bodyweight daily for 5 days prior to subcutaneous tumor injections, after which a half-dose of tamoxifen (50 mg/kg bodyweight) was i.p. injected every 3 days to maintain knockout efficiency. Both genotypes (*LysM^CreERT2^*.*Ube2n^fl/fl^* and *LysM^WT^.Ube2n^fl/fl^* littermate controls) were treated with i.p. tamoxifen to control for the effects of the drug. Mice were euthanized at the indicated time point, and tissue samples were collected after euthanasia by CO^2^ inhalation and decapitation.

### Cell culture

All culture media and supplements were purchased from Thermo Fisher Scientific. The YUMM1.7 murine melanoma cell line was obtained from ATCC via the Duke Cell Culture Facility. *Rosa26^CreERT2^*.*Ube2n^C87S^* murine bone marrow and the UC-764865 compound were provided by Dr. Daniel Starczynowski (Cincinnati Children’s Hospital Medical Center, Cincinnati, Ohio, USA). Yumm1.7 cells were cultured in DMEM: F12 supplemented with 10% FBS and 1× Antibiotic-Antimycotic. Murine bone marrow-derived macrophages (BMDMs) were cultured in RPMI-1640 supplemented with 10% FBS, 10 ng/mL GM-CSF, and 1x Antibiotic-Antimycotic. Cells were cultured in a 37°C incubator supplemented with 5% CO2. All cell lines were tested and confirmed to be free of mycoplasma contamination.

### Immunofluorescence and image analysis

Tumors were frozen in OCT and 12-25µm cryosections were used for immunofluorescence staining. Sections were fixed in methanol for 15 min at −20°C. After washing with PBS, sections were blocked with 10% donkey serum in 0.05% Tween-20 PBS for 30 minutes at room temperature. Sections were incubated with primary antibodies diluted in 10% donkey serum + 0.05% Tween-20 + PBS at optimized concentrations, with incubation overnight at 4°C, followed by incubation with fluorophore-conjugated secondary antibodies diluted to optimal concentrations. Secondary antibodies were purchased from Thermo Fisher Scientific. Images were taken using an Olympus IX73 microscope (Olympus, Tokyo, Japan) with a UPLSAPO 10× or 20× objective. Image analyses were performed on randomly selected fields. Immunofluorescent image acquisition and quantitative analysis were performed using CellSens (version 2.3; Olympus Corporation, Tokyo, Japan). Colocalization analysis (Manders coefficient) was performed using the Colocalization module in CellSens.

### Murine bone marrow extraction

Bone marrow was extracted from the hind limbs of *LysM^CreERT2^Ube2n^fl/fl^*mice and littermate controls as described previously [28]. Briefly, mice were euthanized by CO2 inhalation followed by decapitation. The hind limbs were sprayed with 70% ethanol and dissected to expose the femur and tibia. A 23G needle filled with complete RPMI-1640 was used to flush the bone marrow from the extracted bones. Extracted bone marrow cells were filtered through a 70-micron filter before culturing in differentiation media (RPMI-1640 + 10% FBS + 10 ng/mL M-CSF).

### Collection of BMDM and melanoma tumor cell conditioned media

Differentiated *LysM^CreERT2^*.*Ube2n^fl/fl^*mice and littermate control BMDMs were treated with 1.5 uM 4-OHT for 72 hours, then stimulated with 30% v/v YUMM1.7-conditioned media overnight. All media was then removed from the BMDMs and they were rinsed with sterile PBS twice. Fresh, complete RPMI-1640 (5% FBS) was then added and allowed to incubate for 24 hours. BMDM-CM was then collected, passed through a 40-micron cell strainer, and centrifuged at 20,000 rpm for 5 minutes to remove and pellet any cell debris. Yumm1.7-conditioned media (CM) was generated by growing Yumm1.7 cells to 90% confluence in complete DMEM-F12, then replacing the media with serum-free RPMI-1640 + 1x A-A and culturing overnight. YUMM1.7-CM was gently removed from the cells, passed through a 40-micron cell strainer, then centrifuged at 20,000 rpm for 5 minutes to pellet and remove any cell debris.

### Real-time quantitative PCR

Real-time quantitative PCR was performed with total RNA extracted from cells using TRIzol reagent. RNA (0.1-1 µg) was quantified, and a standardized amount of RNA was reverse transcribed into cDNA using the iScript Reverse Transcription Supermix System. The thermal-cycler program was 25°C for 5 min, 46°C for 20 min, 95°C for 1 min with a permanent hold at 4°C. cDNA was diluted 1:5 before qPCR which was performed on an Applied Biosystems StepOne Real-Time PCR System using selected primers and 2× Universal SYBR Green Fast qPCR Mix. The real-time PCR program was 95°C for 15 min, followed by 40 cycles of 94°C for 20 s, 59°C for 30 s, and 72°C for 30 s. Cycle threshold (Ct) values were normalized to housekeeping genes *Gapdh* and *Stx5a* as an internal control (Table S1).

### ELISA

Enzyme-linked immunoabsorbance assay (ELISA) was used to quantify SPP1 in BMDM-CM using a commercial ELISA kit (RayBiotech, Peachtree Corners, GA, USA) according to the manufacturer’s instructions. Briefly, BMDM-CM were collected following differentiation and induction, centrifuged to remove cellular debris, as described above, and stored at −80 °C until analysis. Standards and samples were added to 96-well microplates pre-coated with an anti-mouse SPP1capture antibody and incubated to allow antigen binding. After washing to remove unbound material, a biotinylated detection antibody specific for mouse SPP1 was added, followed by incubation with HRP-conjugated streptavidin. Signal was developed using tetramethylbenzidine substrate, and the reaction was terminated with stop solution. Absorbance was measured at 450 nm using a microplate reader, and protein concentrations were calculated from a standard curve generated using recombinant mouse SPP1 standards provided with the kit. Concentrations were interpolated from the standard curve using four-parameter logistic regression.

### Flow cytometry

Cells were harvested from their respective culture conditions, washed twice with 1× PBS supplemented with 2% heat-inactivated fetal bovine serum (HI-FBS), and incubated with an anti-CD16/CD32 Fc receptor (FcR) blocking antibody for 10 min at 4°C. Fluorochrome-conjugated antibodies against the indicated surface markers were then added directly to the blocked cells at a 1:200 dilution and incubated for 30 min at 4°C in the dark. Dead cells were also excluded using a L/D viability dye (L/D Ghost 510) at a 1:1000 dilution in serum-free 1× PBS for 20 min at 4°C in the dark. Following staining, cells were washed twice and resuspended in 1× FACS buffer for analysis. Data were acquired on a BD LSRFortessa X-20 flow cytometer, and compensation was performed using single-stained controls prepared from mouse spleen cells. Doublets were excluded based on forward scatter area (FSC-A) versus forward scatter height (FSC-H) parameters, and live cells were subsequently gated for analysis of T cell marker expression. Flow cytometry data were analyzed using FlowJo software (v10; BD Biosciences).

### Pan T-cell isolation and culture

Splenocytes from C57BL/6 chilled mouse spleens were gently isolated via 1-2 mm^3^ sectioning and mechanical dissociation on a 70um filter. Cells were filtered and rinsed into pre-chilled 1xPBS containing DNase-I (100 ug/mL), counted, and purified via a Pan-T-cell negative isolation kit per manufacturer protocol. Pan-T cells were then re-plated at 1.0e^6^ cells/mL into a 96-well flat-bottom TC plate in 0.2 mL of pre-warmed T cell media (RPMI-1640 + 10% HI-FBS + 2mM L-Glutamine + 50 uM 2-ME + 1X NEAA + 10 ng/mL mIL-2 + 100U/mL P/S). T cells were activated with plate-bound anti-CD3 mAb (5 ug/mL; 100 uL for 2h/37C; 145-2C11) and soluble anti-CD28 mAb (2 ug/mL; 37.51) for 72h. T cells were also activated in parallel via 2:1 anti-CD3/CD28 Dynabeads in a 96-well U-Bottom suspension plate. After 72h, activated Pan-T cells from both conditions were harvested, 1X D-PBS washed, and re-plated separately at 1.0e^6^ cells/mL in 0.2 mL T cell media in fresh 96-well flat-bottom TC plates pre-treated with anti-CD3 mAb (5 ug/mL; 100 uL for 2h/37C; 145-2C11).

### T-cell checkpoint expression in co-culture with tumor conditioned BMDM

*LysM^WT^.Ube2n^fl/fl^* and *LysM^CreER^.Ube2n^fl/fl^* bone marrow cells were isolated from n=4-6 mice per group as described above, plated into 1.0 mL BMDM differentiation media in 12-well TC plates at 1.0e^6^ cells/well (RPMI-1640 + 10% HI-FBS + 2mM L-Glutamine + 10 ng/mL GM-CSF + 1000U/mL P/S + 1X A-A), and differentiated for 6 days at humidified 37C, 5% CO_2_. At 3 days after differentiation, BMDMs were treated with 1.5 uM 4-OHT for 3 days in differentiation media. During the final day of differentiation, 30% (v/v) YUMM1.7 tumor cell-CM (TCM) was applied to BMDMs to generate TAM mimics. Post-differentiation and conditioning, TAMs were washed with 1X D-PBS, lifted with 0.5 mM EDTA, and re-plated into 96-well flat-bottom TC plates.

Pan-T cells harvested from matched mouse spleens 6 days prior to co-culture via mechanical tissue disruption and isolation with Pan-T-cell negative isolation kit as described above. Cells were stimulated for 48h with plate-bound anti-CD3e (3 ug/mL, 500 uL; 2h/37C) and soluble anti-CD28 (1 ug/mL) in a 24-well flat-bottom plate at 1.0e^6^ cells/mL in 1.0 mL T-cell media. After 48 hours stimulation, Pan T-cells were harvested via 1xDPBS + ice-cold 10 mM EDTA treatment and transferred onto fresh anti-CD3 mAb coated plates (1 ug/mL, 500 uL; 2h/37C) in 1.0 mL of fresh T-cell media with 30% (v/v) YUMM1.7 TCM at 1.0e^6^ cells/mL and the process was repeated every 48h. After 6 days of activation and conditioning as described above, Pan T-cells were harvested via 1x DPBS + ice-cold 10 mM EDTA treatment, 1X D-PBS washed, and re-plated in T cell media directly on top of TAMs for co-culture. Final co-culture wells contained 0.2 mL of media at a 1:1 T:M ratio (50k:50k). Some wells received 10.0 ug/mL *α*-SPP1 neutralizing antibody (BioXcell) or 10 ug/mL IgG control (BioXcell) upon co-culture. After 4 days of co-culture, cells were harvested for flow cytometry analysis.

### T-cell response to chronic antigen stimulation and BMDM-CM

To mimic chronic antigen-induced T-cell exhaustion, cells received 30% conditioned media from 24h *LysM^WT^.Ube2n^fl/fl^* and *LysM^CreER^.Ube2n^fl/fl^* BMDMs stimulated with Yumm1.7 cell conditioned media. Every 72h, T cells were passaged into freshly prepared wells with 50% media renewal. At baseline, day 3, and day 9 of chronic *in vitro* stimulation, T cells were stained and fixed for flow cytometry analysis.

### Statistical analysis

Statistical analysis was performed using GraphPad Prism (version 9.0.0, La Jolla, California). All data are reported as means +/- SD unless otherwise stated. Comparisons between groups were performed using an unpaired two-tailed t-test with Welch’s correction, which does not assume equal variance. Differences in means across groups with repeated measures over time were analyzed using two-way repeated-measures ANOVA with Bonferroni’s multiple comparisons post-hoc test. Analyses with p<0.05 were considered statistically significant.

## RESULTS

### Conditionally induced U*be2n* deletion specifically in myeloid cells hinders YUMM1.7 tumor growth

We asked whether *Ube2n* deletion in myeloid cells decreases tumor growth. To answer this question, we employed the *LysM^CreER^.Ube2n^fl/fl^* mouse model to enable a temporally inducible myeloid-specific deletion of *Ube2n (Ube2n^MyeKO^)*. We administered i.p. injections of tamoxifen (100mg/kg bodyweight) to *LysM^CreER^.Ube2n^fl/fl^* mice and littermate controls daily for 5 days (Fig. 1A). We then subcutaneously injected YUMM1.7 cells into the back skin and administered tamoxifen (50mg/kg) every three days. Tumor growth and body weight were recorded, and tumors were collected at the endpoint. YUMM1.7 tumors displayed significantly decreased growth over time, culminating in reduced tumor weights at endpoint in *Ube2n^MyeKO^*animals (Fig. 1B-D). These results indicate that *Ube2n* loss in myeloid cells hinders YUMM1.7 growth.

**Figure 1:**
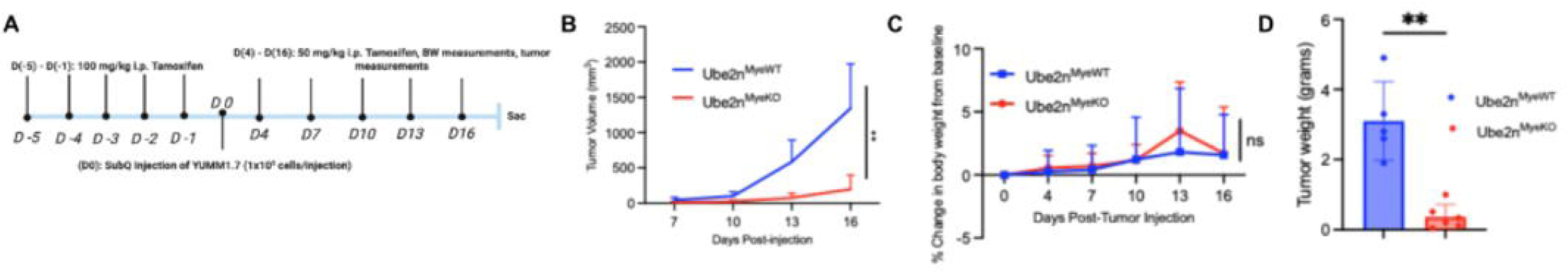
Myeloid-specific *Ube2n* deletion hinders YUMM1.7 melanoma growth. (A) Experimental strategy. Mice received 100 mg/kg i.p. tamoxifen daily on days –5 to –1, followed by subcutaneous injection of 1 x 10^5^ YUMM1.7 cells on day 0. 50 mg/kg i.p. tamoxifen was administered every 3 days on days 4-16. Body weight was recorded on days 0-16. Tumors were measured on days 7-16. (B) Quantification of YUMM1.7 tumor growth kinetics, mean ± SD. n=6 *Ube2n^MyeKO,^*n=5 *Ube2n^MyeWT^*. Analyzed by two-way repeated measures ANOVA with Bonferroni post-hoc test performed in GraphPad Prism. (C) Percent change in body weight from day 0, mean +/- SD. n=6 *Ube2n^MyeKO,^* n=5 *Ube2n^MyeWT^.* Analyzed by two-way repeated-measures ANOVA with Bonferroni’s multiple comparisons post-hoc test. Body weight did not differ between genotypes and there was no genotype x time interaction. Bonferroni-corrected comparisons between genotypes was non-significant at every time point. All group means represented net weight gain relative to baseline. (D) Comparison of final tumor weights presented as mean with individual values +/- SD. n=6 *Ube2n^MyeKO^*, n=5 *Ube2n^MyeWT^.* Statistical significance was determined using an unpaired two-tailed t-test with Welch’s correction. *p < 0.05, **p < 0.01, ***p < 0.001. ns= not significant.

### U*be2n* deletion decreases myeloid cell burden and alters T cell checkpoint expression

To investigate alterations in the immune landscape of the *Ube2n^MyeKO^*tumors, we used fluorescent immunostaining (IF) to identify and quantify myeloid cells. We looked for myeloid-specific knockout of *Ube2n* by co-staining OCT-embedded tissues for myeloid cells (CD11B) and UBE2N (Fig. 2A). Co-localization analysis showed a significant decrease in the fraction of CD11B+ signal co-occupied by UBE2N (Fig. 2B). We also observed a significant decrease in CD11B+ signal in *Ube2n^MyeKO^* tumors (Fig. 2C). We questioned whether checkpoint expression was altered in the TME of *Ube2n^MyeKO^* tumors. Staining for PD-1 and its ligand PD-L1 revealed a decrease in the fraction of CD3+ signal co-occupied by PD-1 in *Ube2n^MyeKO^* tumors (Fig. 2 D-E). This was accompanied by a generalized decrease in PD-L1 expression in the TME of *Ube2n^MyeKO^* tumors compared to *Ube2n*^MyeWT^ controls (Fig. 2 F-G). These findings suggest that *Ube2n* loss in myeloid cells decreases myeloid cell burden in the tumor microenvironment and alters the expression of checkpoint markers in the TME.

**Figure 2:**
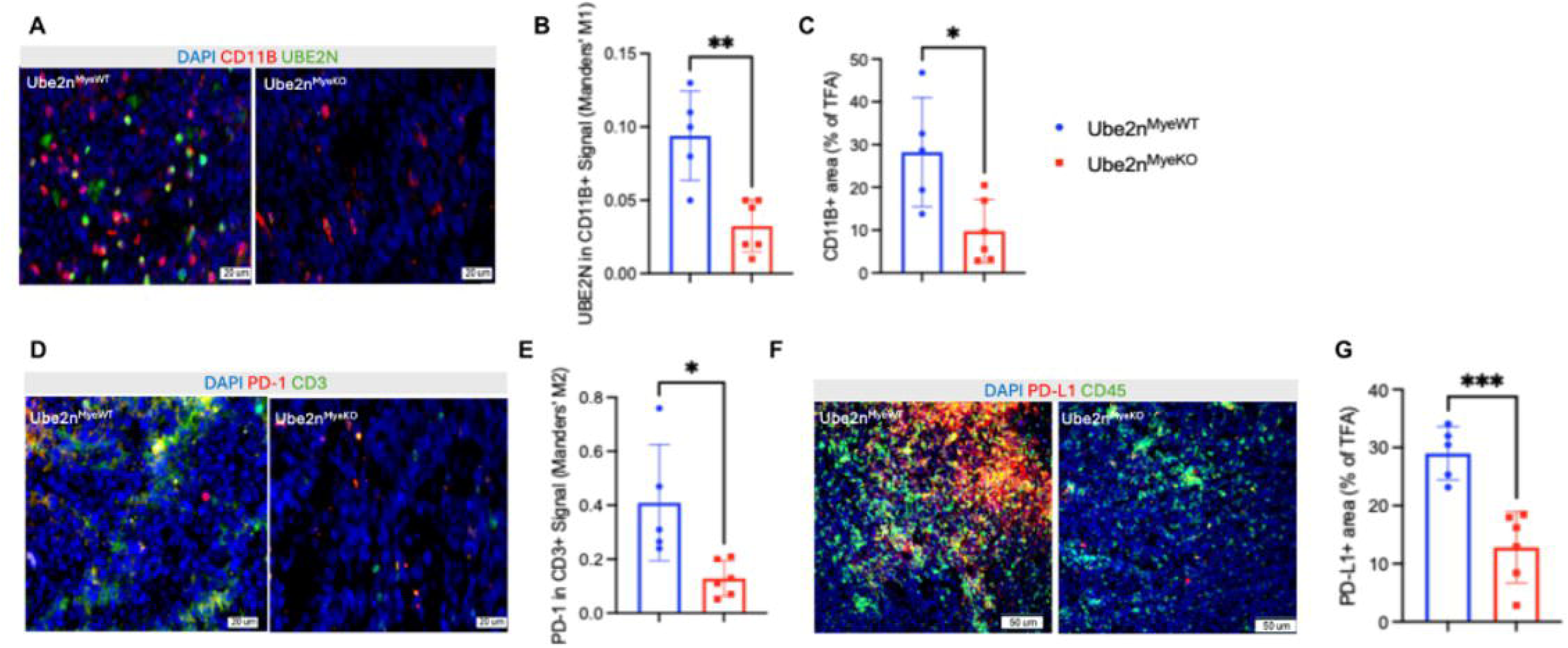
Myeloid *Ube2n* deletion decreases intratumoral myeloid area and checkpoint marker abundance in YUMM1.7 tumors. (A) Representative immunofluorescence images of tumor sections from *Ube2n^MyeWT^* and *Ube2n^MyeKO^* animals stained for the myeloid cell marker CD11B (red), and UBE2N (green) with DAPI nuclear counterstain (blue). Scale bar = 20 um (B) Quantification of UBE2N within the CD11B+ compartment, expressed as Manders’ coefficient M1. M1 represents the fraction of CD11B+ signal co-occupied by UBE2N. (C) Quantification of CD11B+ area normalized to total fluorescent area (TFA). TFA = the summed area of all fluorescent signal within the imaged field. (D) Representative images stained for PD-1 (red) and CD3 (green) with DAPI nuclear counterstain (blue). Scale bar = 20 um. (E) Quantification of PD-1 within the T cell compartment, expressed as Manders’ coefficient M2. M2 represents the fraction of CD3+ signal co-occupied by PD-1. (F) Representative images stained for PD-L1 (red) and pan-leukocyte marker CD45 (green) with DAPI nuclear counterstain (blue). Scale bar=50 um. (G) Quantification of PD-L1+ area normalized to TFA. Field-level measurements were averaged within each tumor, and all statistical comparisons were performed at the level of the animal. Each point represents one tumor. Data are shown as mean +/- SD, n=6 *Ube2n^MyeKO^* and n=5 *Ube2n^MyeWT^* tumors. Colocalization analysis was performed in CellSens using a manual threshold applied identically to all images and across both genotypes. Statistical significance was determined by unpaired two-tailed t-test with Welch’s correction. *p < 0.05, **p < 0.01, ***p < 0.001.

### U*be2n* deletion in macrophages alters expression of inflammatory and immunosuppresive mediators

We questioned whether phenotypic changes occur in *Ube2n^MyeKO^*macrophages. To investigate this, we collected bone marrow from *LysM^CreER^Ube2n^fl/fl^* mice and littermate controls to generate bone marrow-derived macrophages (BMDMs) *in vitro*. BMDMs were differentiated, treated with 4-OHT for 72 hours to induce *Ube2n*-knockout (KO), then stimulated with YUMM1.7 tumor-conditioned media (TCM) for 12 hours. Cells were then collected for isolation of total RNA and RT-qPCR analysis. We first validated the *Ube2n*-KO in *Ube2n^MyeKO^* group (Fig. 3A). Reduction of UBE2N at the protein level was confirmed by Western blot of 4-OHT-treated *Ube2n^MyeKO^*and *Ube2n^MyeWT^* BMDM lysates (Fig. S3). In addition, we found that a range of genetic markers previously associated with both pro- and anti-inflammatory macrophage identity and function were decreased in *Ube2n^MyeKO^* BMDMs. These include *Spp1, Tnfa*, *Apoe, and Vegfa* (Fig 3B-E).

**Figure 3:**
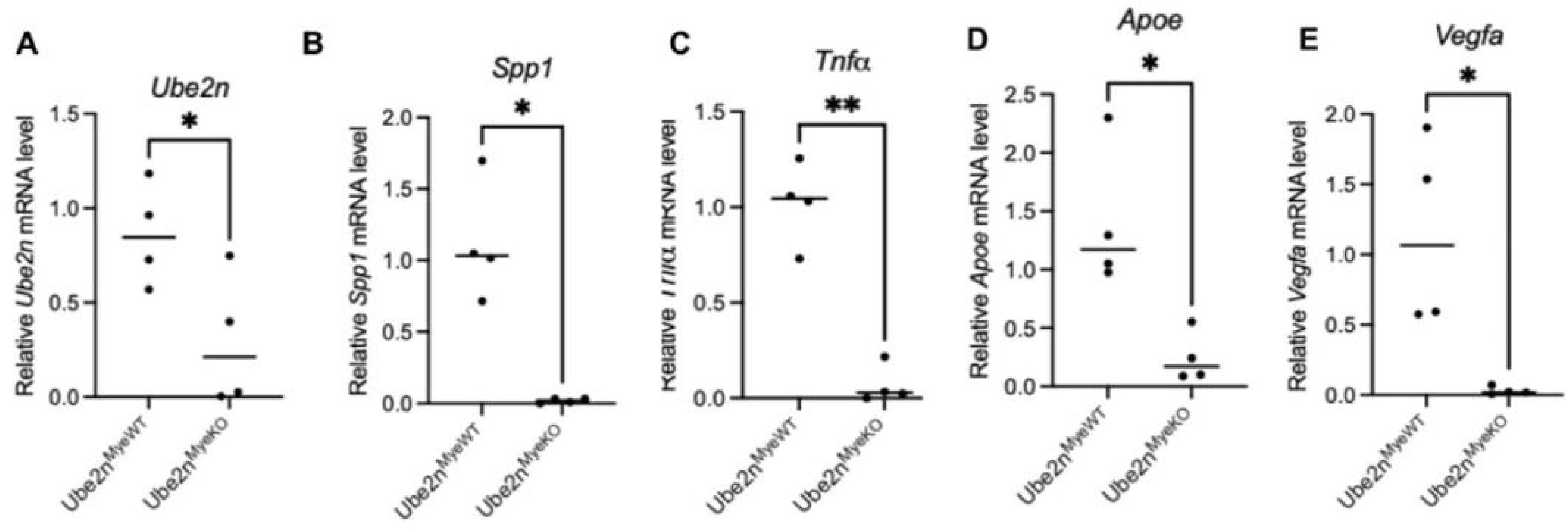
UBE2N deletion in macrophages decreases Ube2n-dependent inflammatory and immunosuppressive genes. (A-E) Relative mRNA expression of *Ube2n*, *Spp1*, *Tnfa, Apoe,* and *Vegfa* in either *Ube2n^MyeKO^*or *Ube2n^MyeWT^* BMDMs treated with 4-OHT for 72 hours then stimulated with TCM for 12 hours. Each point represents one independent bone marrow preparation. Technical replicates were averaged within each preparation before analysis. Data are shown as means with individual values. N= 4 independent bone marrow preparations per genotype. All data are normalized to the *Gapdh* and *Stx5a* housekeeping genes. Statistical significance was determined using an unpaired two-tailed t-test with Welch’s correction. *p < 0.05, **p < 0.01, ***p < 0.001.

### SPP1 is decreased in *Ube2n^MyeKO^* tumors and bone marrow-derived macrophages

Because SPP1 is downregulated in our *Ube2n^MyeKO^* BMDMs and has a well-defined role in promoting tumor progression via expression in TAMs, we asked whether SPP1 could be an effector of myeloid UBE2N. Fluorescent immunostaining showed a generalized decrease of SPP1 in the TME of *Ube2n^MyeKO^* tumors (Fig. 4A-B). To probe for a possible link between UBE2N and SPP1, we used BMDMs from *Rosa^CreER^Ube2n^fl/fl.C87S^* mice, which harbor a tamoxifen-inducible knock-in mutation that reduces UBE2N catalytic function. *Rosa^CreER^Ube2n^fl/fl.C87S^* (Ube2n^C87S.KI^) and *Rosa^CreER^Ube2n^WT^* (Ube2n^WT^) BMDMs were polarized toward either M1-like or M2-like phenotypes and treated with 4-OHT and then collected for RNA extraction. RT-qPCR revealed a significantly elevated level of *Spp1* expression in M2-polarized BMDMs relative to M1. *Ube2n*^C87S.KI^ resulted in a significant decrease in *Spp1* in both M1- and M2-polarized macrophages (Fig. 4C). We also observed that *Ube2n* deletion in BMDMs decreased SPP1 secretion via enzyme-linked immunosorbance assay (Fig. 4D). To explore a possible link between macrophage-derived SPP1 and T cell checkpoint marker expression, we cultured chronically activated pan T-cells in the presence of conditioned media (CM) collected from either *Ube2n^MyeWT^* or *Ube2n^MyeKO^* BMDMs stimulated with YUMM1.7 tumor cell-conditioned media (TCM) (Fig. 4E). After 9 days in culture, chronically activated T-cells were then collected and expression of checkpoint markers PD-1, TIM-3 and LAG-3 was quantified using flow cytometry. We observed that the percentages of PD-1+ TIM-3+ LAG-3+ triple positive cells over total CD8+ T-cells were significantly higher in the *Ube2n^MyeWT^* BMDM-CM-treated group than *Ube2n^MyeKO^*BMDM-CM-treated group (Fig. 4F). The gating strategy for this analysis is shown in (Fig. S1). We then asked whether SPP1 could be neutralized in *Ube2n^MyeWT^* BMDM-CM to mitigate expression of the checkpoint marker PD-1. Treatment with an anti-SPP1 neutralizing antibody significantly decreased PD-1 expression in activated T-cells cultures in the presence of *Ube2n^MyeWT^* BMDM-CM (Fig. 4G). The gating strategy for this is shown in (Fig. S2). This observation suggests that SPP1 is more abundant in *Ube2n^MyeWT^*BMDM-CM and can be neutralized to mitigate checkpoint marker expression on chronically activated T cells. Collectively, these data indicate that myeloid UBE2N orchestrates a distinct secretory program in BMDMs. SPP1 is identified here as a candidate effector that may play a role in driving checkpoint marker expression in the YUMM1.7 TME and on chronically activated T cells.

**Figure 4:**
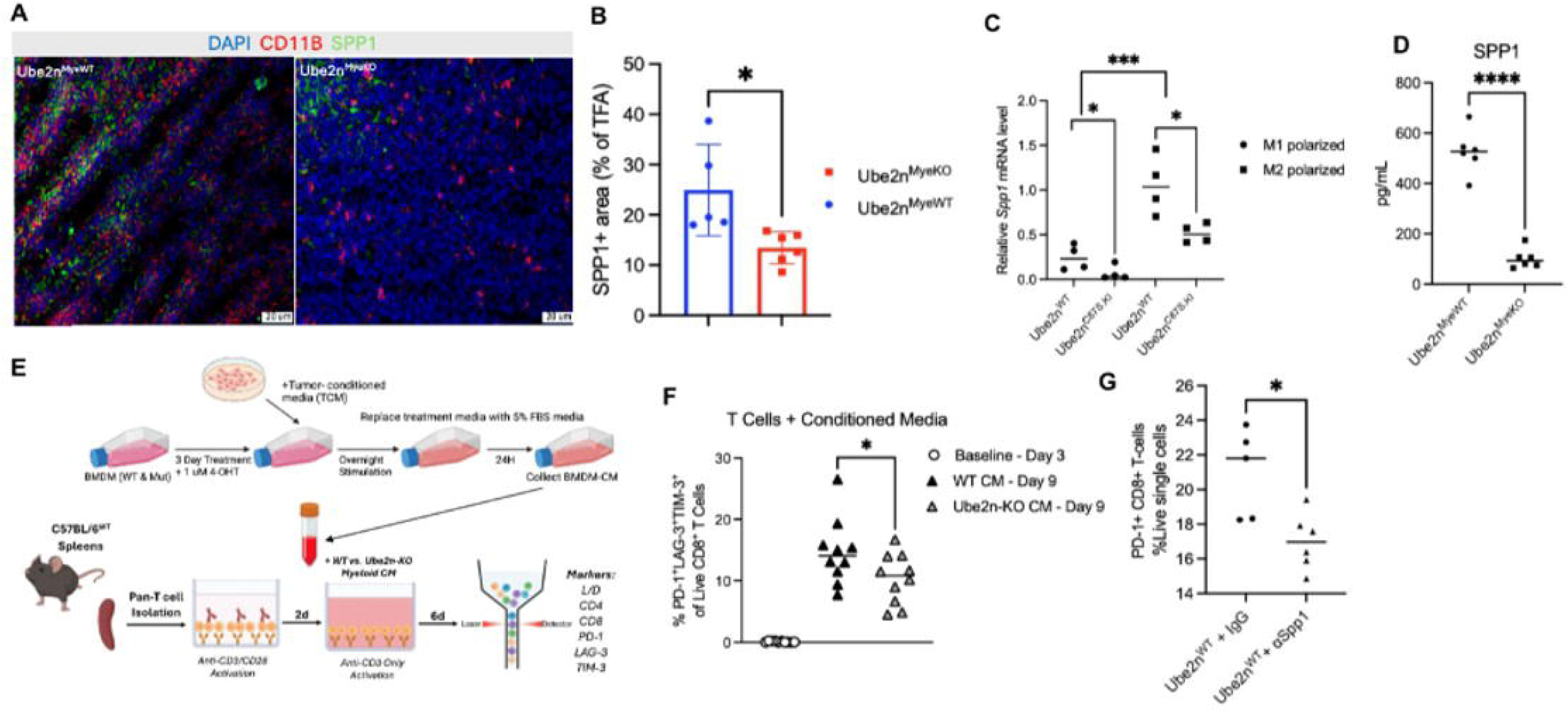
Myeloid *Ube2n* supports SPP1 production, and myeloid-derived SPP1 alters CD8+ T cell checkpoint marker expression. (A) Representative immunofluorescence images of tumor sections from *Ube2n^MyeKO^* and *Ube2n^MyeWT^* animals stained for CD11B (red), SPP1 (green), and DAPI nuclear counterstain (blue). Scale bar = 20 um. (B) Quantification of SPP1+ area normalized to TFA. Each point represents one tumor. Field-level measurements were averaged within each tumor. Comparisons were performed at the level of the animal. (C) Relative mRNA level of Spp1 in M1- and M2-polarized RosaCreERT2.Ube2nC87S.KI and RosaCreERT2.Ube2nWT BMDMs treated with 4-OHT. Each point represents an independent culture. N=4 cultures per condition. Data are shown as means with individual values +/- SD. Statistical significance was determined using an unpaired two-tailed t-test with Welch’s correction. (D) SPP1 protein concentration in BMDM culture media supernatant from 4-OHT treated Ube2nMyeKO and Ube2nMyeWT BMDMs. Each point represents an independent culture. N=6 cultures per genotype. (E) Scheme of T-cell:BMDM-CM experiment. (F) Frequency of PD-1+ LAG-3+ TIM-3+ cells among live CD8+ T cells at baseline (day 3) and after 6 days of culture (day 9) in either *Ube2n^MyeKO^* or *Ube2n^MyeWT^* BMDM-CM. Each point represents an individual culture well. (G) Frequency of PD-1+ CD8+ T cells among live single cells following co-culture with *Ube2n^MyeWT^* BMDMs in the presence of an IgG isotype control or SPP1-neutralizing antibody (aSPP1). Each point represents an individual culture well. Data are shown as individual values + SD. Statistical significance was determined using an unpaired two-tailed t-test with Welch’s correction. *p < 0.05, **p < 0.01, ***p < 0.001, ****p<0.0001.

## DISCUSSION

In the setting of myeloid-specific *Ube2n* deletion, we observed a decreased YUMM1.7 tumor growth phenotype. Our in vivo results indicate that myeloid cells may be depleted in the YUMM1.7 TME by myeloid-specific deletion of Ube2n. We observed decreased checkpoint marker expression in the *Ube2n^MyeKO^* TME. In vivo and in vitro analyses showed a consistent decrease in SPP1 expression and secretion in the context of myeloid-specific Ube2n knockout. It is well-documented that myeloid cells, particularly TAMs, may either enhance or inhibit T-cell activity in the TME [30,31,32]. Our analysis of the effects of *Ube2n^MyeKO^*BMDM-CM suggest that Ube2n-deficient BMDMs may alter checkpoint expression on chronically activated T cells.

Abundance of myeloid cells, specifically immunosuppressive TAM, has consistently been correlated with poor prognosis and aggressive tumor behavior across a variety of cancers [33,34,35,36]. Given this clinical relevance, it is necessary to determine whether myeloid cells in the TME may be reduced, reprogrammed to execute anti-tumor immune functions, or both. Our findings suggest that myeloid cell reduction may be accomplished in the YUMM1.7 TME.

Markers and functional qualities of pro-tumor macrophages are documented extensively in the literature, including SPP1 [37,38,39]. Our profiling of *Ube2n*^MyeKO^ macrophages demonstrated a decrease in expression and secretion of Spp1. Recent single-cell transcriptomic studies have identified SPP1 as a conserved marker of immunosuppressive TAMs across multiple malignancies, including colorectal, breast, lung, and liver cancers, where SPP1+ TAMs promote extracellular matrix remodeling, angiogenesis, and T-cell dysfunction [39,40,41,42,43]. Spp1 expression in TAMs has been directly linked to T-cell exhaustion in the TME [22,23,44]. Our findings suggest that myeloid-specific knockout of Ube2n decreases SPP1 in the YUMM1.7 TME. Our profiling of *Ube2n^MyeKO^* BMDMs suggests that Spp1 expression and secretion are decreased in Ube2n-deficient BMDMs and have an indirect effect on checkpoint marker expression on T cells. Blocking SPP1 with a neutralizing antibody dampens WT BMDM-induced PD-1 checkpoint marker expression on CD8+ T cells. These findings suggest that Spp1 may be a downstream effector of UBE2N in myeloid cells.

Although a distinct regulatory relationship between Ube2n and Spp1 has not been fully elucidated, several lines of evidence suggest that UBE2N influences Spp1 expression through its role in inflammatory signaling. UBE2N catalyzes K63 ubiquitination, a post-translational modification required for activation of TRAF6/TAK1/NF-κB signaling, an established regulator of innate immunity [45, 46, 47]. Given that SPP1 is a known marker of TAM states and is transcriptionally induced by NF-κB-responsive signaling, it is possible that UBE2N contributes to the emergence of SPP1⁺ macrophages by maintaining pro-inflammatory transcriptional programs. Further experimental validation is required to determine how UBE2N regulates SPP1 expression through NF-κB.

Several limitations of the current study should be acknowledged. Although our data demonstrate alterations in immune cell abundance and checkpoint marker expression following myeloid Ube2n deletion, the precise molecular mechanisms linking UBE2N activity to macrophage recruitment and polarization remain to be fully defined. A limitation of the *LysM^CreER^* model is that all myeloid lineage cells are affected by the knockout [48], not solely macrophages. Deletion efficiency in LysM^CreER^ mice varies from over 90% in granulocytes, 83-89% in mature macrophages, and 16% in dendritic cells [48]. More specific targeting strategies are needed to define cell-type-specific functions. A major limitation imposed by the dosing schedule used in this study is that the Ube2n knockout is induced in advance of the YUMM1.7 injection, and thus the study does not address whether UBE2N can be therapeutically targeted to address melanoma. Additionally, because Vegfa, Tnfa, and Apoe are concurrently reduced in Ube2n-deficient BMDMs, the SPP1-neutralization experiment does not exclude contribution from these or other secreted factors to the observed changes in T cell checkpoint marker expression. SPP1 is therefore best described as one candidate effector among several rather than as a dominant mediator. Finally, it must be noted that PD-L1 checkpoint expression is quantified here without respect to cell type. Further study will be needed to determine whether PD-L1 checkpoint expression is decreased in leukocytes or the cancer and stromal cells.

Taken together, our results demonstrate that UBE2N expression in myeloid cells influences myeloid cell abundance, immune checkpoint expression, and macrophage secretory program. UBE2N may drive expression of SPP1 to exert an indirect effect on T cell checkpoint marker expression. This study suggests the possibility of targeting UBE2N in myeloid cells to combat YUMM1.7 growth and effect changes checkpoint marker expression on T cells and in the YUMM1.7 TME.

## Supporting information

Supplementary Figures and Key Resources

## AUTHOR CONTRIBUTIONS

Conceptualization: JZ, KS

Methodology: KS, AP, KZ, AK, DS

Investigation: KS, AP, JZ

Visualization: KS, AP

Funding acquisition: JZ

Project administration: JZ

Supervision: JZ

Writing – original draft: KS, JZ

Writing – review & editing: KS, JZ, AP, KZ, AK, DS

## DATA AVAILABILITY STATEMENT

All source data are available in the figures and upon request.

## ACKNOWLEDGEMENTS

We thank Dr. Shizuo Akira of Osaka University and Dr. Shao-Cong Sun of University of Texas MD Anderson for providing the *Rosa26^Cre-ERT2^Ube2n^fl/fl^* mice, as well as Fang Li (Duke University, Durham, NC) for sharing resources and expertise in cell culture. We thank Dr. Georgia Beasley of the Duke Medical School Department of Surgery for her guidance and expertise.

