## Supplementary Figures and Key Resources for "Conditional Myeloid-Specific Inhibition of UBE2N Hinders YUMM1.7 Growth"

This file includes:

Figure S1.
Figure S2.
Figure S3.
Table S1. RT-PCR-Primers
Table S2. Key resources table

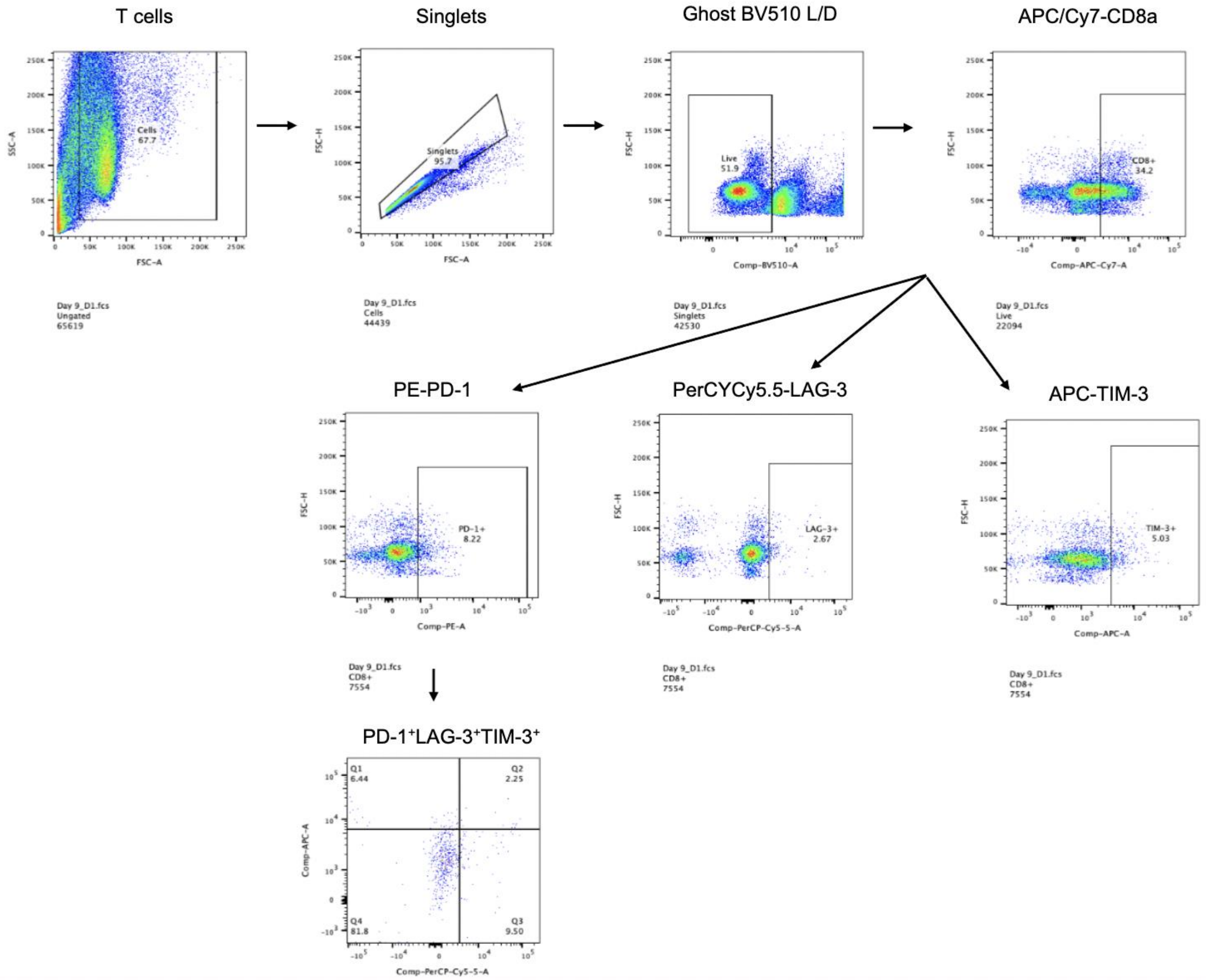

**Supplemental Figure S1: Gating strategy for Fig. 4(F), T-cell/BMDM-CM co-culture.** T-cell population isolated from debris into singlets, live cells, and PD-1+ LAG-3+TIM-3+ live single cells as the final gate.

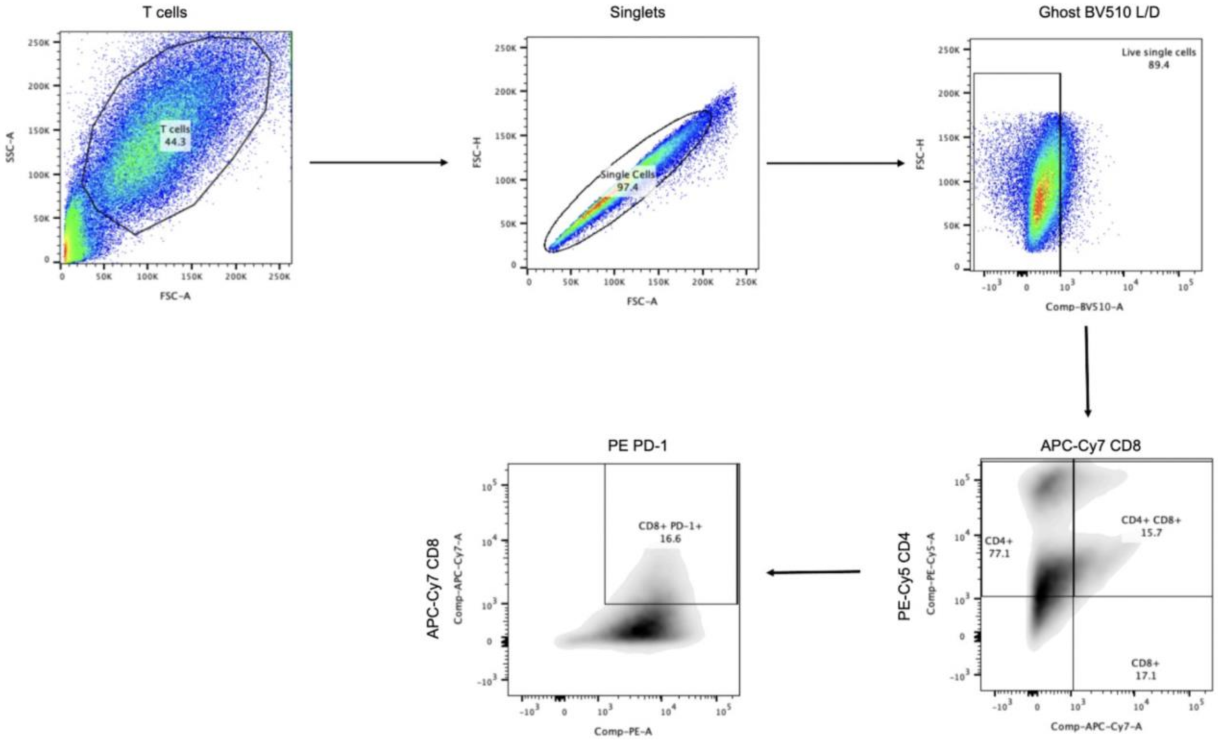

**Supplementary Figure S2: Gating strategy for Fig. 4(G), T-cell/BMDM co-culture.** T-cell population isolated from debris into singlets, live cells, and PD-1+ CD8+ CD4- live single cells as the final gate.

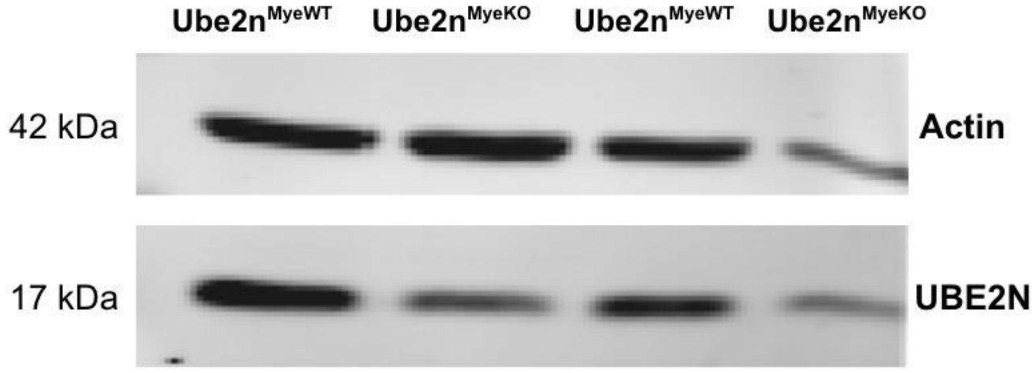

**Supplementary Figure S3: UBE2N protein is reduced in Ube2nMyeKO bone marrow-derived macrophages**. Immunoblot of whole-cell lysates from BMDMs generated from *Ube2n*$^{MyeWT}$ and *Ube2n*$^{MyeKO}$ mice probed for UBE2N (17 kDa) and Actin (42 kDa) as a loading control. Lanes 1-2 and 3-4 represent two independent bone marrow preparations. Actin signal is reduced in lane 4, indicating a lower total protein load in that lane. The load-matched comparison of lanes 1 and 2 therefore provides the clearest assessment of UBE2N reduction. Blots were cropped to the region shown.

**Table S1. RT-PCR primers.**

| | |
|---|---|
| Mouse *Ube2n Forward* | TGAGAGCAACGCCCGTTATTT |
| Mouse *Ube2n Reverse* | GCCATTGGGTATTCTTCTGGAA |
| Mouse *Stx5a Forward* | CGGAAACGCTACGGATCTAAG |
| Mouse *Stx5a Reverse* | CAGGGGACAGAACCTGTGT |
| Mouse *Spp1 Forward* | AGCAAGAAACTCTTCCAAGCAA |
| Mouse *Spp1 Reverse* | CTGTCCTGGTATTGAGGGTGG |
| Mouse *Vegfa Forward* | GAGGTCAAGGCTTTTGAAGGC |
| Mouse *Vegfa Reverse* | CCGAAAAGCTGTCCCACAAAA |
| Mouse *Tnfa Forward* | CCCTCACACTCACAAACCAC |
| Mouse *Tnfa Reverse* | ACAAGGTACAACCCATCGGC |
| Mouse *Apoe Forward* | CTGACAGGATGCCTAGCCGA |
| Mouse *Apoe Reverse* | CCTGCTCAGGTTGTTGCTGG |
| Mouse *Gapdh Forward* | GCACAGTCAAGGCCGAGAAT |
| Mouse *Gapdh Reverse* | GCCTTCTCCATGGTGGTGA |
| Mouse Genotyping: *Ube2n wt Forward* | ACACCTTTAATCCCAGCAGAGGCCTAC |
| Mouse Genotyping: *Ube2n common Reverse* | CTCTGCCCCTCTGCTGATCTTTAACAT |
| Mouse Genotyping: *Ube2n ko Forward* | CTAAAGCGCATGCTCCAGACTGCCTTG |

**Table S2. Key resources table.**

| REAGENT or RESOURCE | SOURCE | IDENTIFIER |
|---|---|---|
| **Antibodies** | | |
| Rat IgG | Bio X Cell | Cat#:BE0089; RRID: AB_1107769 |
| FITC anti-mouse CD45 | BioLegend | Cat#:103108; Clone#30-F11; RRID: AB_312973 |
| PE anti-mouse/human CD11b | BioLegend | Cat#:101208; Clone#M1/70; RRID: AB_312791 |
| PE anti-mouse CD8a | BioLegend | Cat#:100708; Clone#53-6.7; RRID: AB_312747 |
| FITC anti-mouse CD3e | BioLegend | Cat#:152304; Clone#500A2; RRID: AB_2632667 |
| PE anti-mouse CD279 (PD-1) | BioLegend | Cat#:135205; Clone#29F.1A12; RRID: AB_1877232 |
| PE anti-mouse CD274 (B7-H1, PD-L1) | BioLegend | Cat#:124308; Clone#:10F.9G2; RRID: AB_2073556 |
| AlexaFluor 488 Donkey anti-Rabbit IgG (H+L) | Invitrogen | Cat#:A21206 |
| DAPI | Thermo Scientific | Cat#:62248 |
| APC/Cyanine7 anti-mouse CD8 | BioLegend | Cat#:100714; Clone#:53-6.7; RRID: AB_312753 |
| PE-Cy5 anti-mouse CD4 | | Cat#:100514; Clone#:RM4-5; RRID: AB_312717 |
| FITC anti-mouse I-A/I-E | BioLegend | Cat#:107605; Clone#M5/114.15.2; RRID: AB_313320 |
| Anti-UBE2N/Ubc13 | Cell Signaling Technology | Cat#:4919; RRID: AB_2211168 |
| TruStain FcX Rat IgG anti-mouse CD16/CD32 | BioLegend | Cat#:101320; Clone#:93; RRID: AB_1574975 |
| Dynabeads™ Mouse T-Activator CD3/CD28 | Thermo Fisher | Cat#11456D |
| Ultra-LEAF™ Purified anti-mouse CD3ε Antibody | BioLegend | Cat#103164; Clone#:30-F1 |
| PerCP/Cyanine5.5 anti-mouse CD223 (LAG-3) Antibody | BioLegend | Cat#:125211 |
| APC anti-mouse CD366 (Tim-3) Antibody RMT3-23 | BioLegend | Cat#:119705 |
| Anti-Osteopontin SPP1 Rabbit Monoclonal Antibody | Boster | Cat#:M00634; Clone#CEO-19; UniProt ID#P10451 |
| InVivoMAb anti-mouse/human/rat osteopontin (SPP1) | BioXcell | Cat#BE0382 |
| InVivoMAb mouse IgG1 isotype control | BioXcell | Cat#:BE0083 |
| **Chemicals, peptides, and recombinant proteins** | | |
| 4-hydroxytamoxifen | Sigma | Cat#:H7904 |
| DMEM | Thermo Fisher | Cat#:11965092 |

| | | |
|---|---|---|
| FBS | R&D | Cat#:S11150H |
| Antibiotic-Antimycotic | Thermo Fisher | Cat#:15240096 |
| RPMI 1640 | Thermo Fisher | Cat#:11875093 |
| Recombinant Mouse M-CSF Protein | R&D | 416-ML-010 |
| Recombinant murine IL-2 | PeproTech | Cat#:212-12 |
| Recombinant murine IL-4 | PeproTech | Cat#:214-14 |
| Recombinant murine IL-13 | PeproTech | Cat#:210-13 |
| Recombinant murine IFNy | PeproTech | Cat#:315-05 |
| Matrigel | Corning | Cat#:354234 |
| Ghost Dye™ Violet 510 | TONBO | Cat#:13-0870-T500 |
| TRIzol reagent | Invitrogen | Cat#:15596018 |
| 2X Universal SYBR Green Fast qPCR Mix | ABclonal | Cat#:RK21203 |
| **Critical commercial assays** | | |
| Pan T-Cell Isolation Kit II | Miltenyi | Cat#:130-095-130 |
| iScript™ Reverse Transcription Supermix System | Bio-Rad | Cat#:1708840 |
| Mouse OPN ELISA Kit | Ray Biotech | Cat#:ELM-OPN-1 |
| Applied Biosystems™ StepOne™ Real-Time PCR System | Thermo Fisher | Cat#:4376357 |
| **Experimental models: Cell lines** | | |
| Yumm1.7 | ATCC | Cat#:CRL-3362; RRID: CVCL_JK16 |
| **Experimental models: Organisms/strains** | | |
| Ube2n$^{C87S}$ Mutant and WT bone marrow | Provided by Dr. Daniel Starczynowski | Cincinnati Children's Hospital Medical Center, Cincinnati, OH, USA |
| Mouse: $LysM^{CreER}$ | Jackson Laboratories | Strain # 032291 |
| $Rosa^{CreER}Ube2n^{fl/fl}$ | Provided by Dr. Shao-Cong Sun | Chinese Institute for Immunology, Chinese Institutes for Medical Research, Beijing, China |
| **Software and algorithms** | | |
| ImageJ/Fiji | Schneider et al.; Schindelin et al. | https://imagej.nih.gov/ij/; RRID: SCR_003070 |
| FlowJo version 10.0 | | https://www.flowjo.com/solutions/flowjo; RRID: SCR_008520 |
| GraphPad Prism version 9.0.0 | | RRID: SCR_002798 |
