## Supplementary material for "Conditional Myeloid-Specific Inhibition of UBE2N Hinders YUMM1.7 Growth": Figures and Legends

**Figures and Legends for:**

**1 Department of Dermatology, Duke University Medical Center, Durham, NC, USA**

**2 Cincinnati Children's Hospital Medical Center, Cincinnati, OH, USA**

**3 Department of Pathology, Duke University Medical Center, Durham, NC, USA**

**This file includes:**

**Figure 1.**

**Figure 2.**

**Figure 3.**

**Figure 4.**

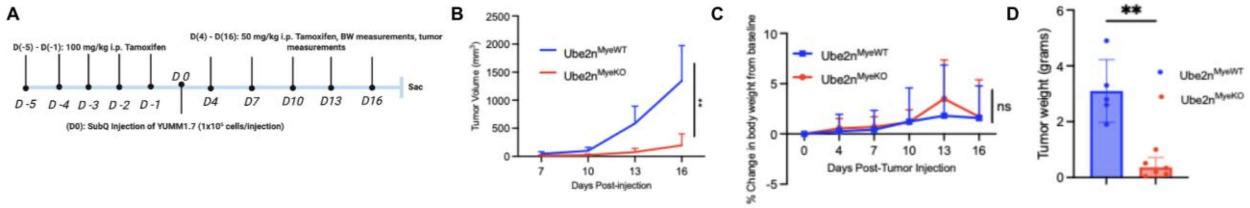


**Figure 1: Myeloid-specific *Ube2n* deletion hinders YUMM1.7 melanoma growth.** (A) Experimental strategy. Mice received 100 mg/kg i.p. tamoxifen daily on days –5 to –1, followed by subcutaneous injection of 1 x $10^5$ YUMM1.7 cells on day 0. 50 mg/kg i.p. tamoxifen was administered every 3 days on days 4-16. Body weight was recorded on days 0-16. Tumors were measured on days 7-16. (B) Quantification of YUMM1.7 tumor growth kinetics, mean ± SD. n=6 *Ube2n*$^{MyeKO,}$ n=5 *Ube2n*$^{MyeWT}$ . Analyzed by two-way repeated measures ANOVA with Bonferroni post-hoc test performed in GraphPad Prism. (C) Percent change in body weight from day 0, mean +/- SD. n=6 *Ube2n*$^{MyeKO,}$ n=5 *Ube2n*$^{MyeWT}$. Analyzed by two-way repeated-measures ANOVA with Bonferroni's multiple comparisons post-hoc test. Body weight did not differ between genotypes and there was no genotype x time interaction. Bonferroni-corrected comparisons between genotypes was non-significant at every time point. All group means represented net weight gain relative to baseline. (D) Comparison of final tumor weights presented as mean with individual values +/- SD. n=6 *Ube2n*$^{MyeKO,}$ n=5 *Ube2n*$^{MyeWT}$. Statistical significance was determined using an unpaired two-tailed t-test with Welch's correction. *p < 0.05, **p < 0.01, ***p < 0.001. ns= not significant.

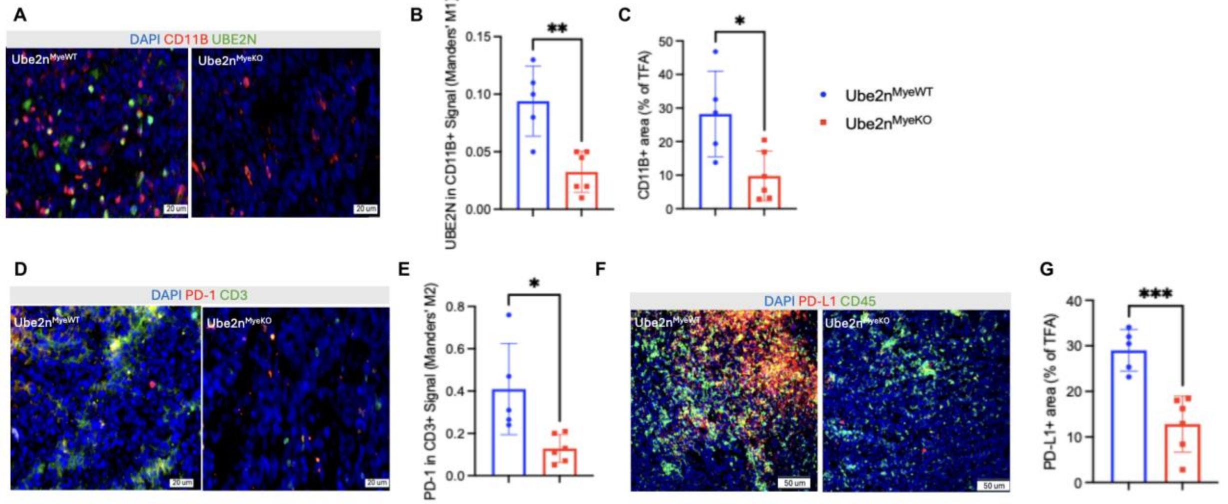


**Figure 2: Myeloid *Ube2n* deletion decreases intratumoral myeloid area and checkpoint marker abundance in YUMM1.7 tumors.** (A) Representative immunofluorescence images of tumor sections from *Ube2n*$^{MyeWT}$ and *Ube2n*$^{MyeKO}$ animals stained for the myeloid cell marker CD11B (red), and UBE2N (green) with DAPI nuclear counterstain (blue). Scale bar = 20 um. (B) Quantification of UBE2N within the CD11B+ compartment, expressed as Manders' coefficient

M1. M1 represents the fraction of CD11B+ signal co-occupied by UBE2N. (C) Quantification of CD11B+ area normalized to total fluorescent area (TFA). TFA = the summed area of all fluorescent signal within the imaged field. (D) Representative images stained for PD-1 (red) and CD3 (green) with DAPI nuclear counterstain (blue). Scale bar = 20 um. (E) Quantification of PD-1 within the T cell compartment, expressed as Manders' coefficient M2. M2 represents the fraction of CD3+ signal co-occupied by PD-1. (F) Representative images stained for PD-L1 (red) and pan-leukocyte marker CD45 (green) with DAPI nuclear counterstain (blue). Scale bar=50 um. (G) Quantification of PD-L1+ area normalized to TFA. Field-level measurements were averaged within each tumor, and all statistical comparisons were performed at the level of the animal. Each point represents one tumor. Data are shown as mean +/- SD, n=6 $Ube2n^{MyeKO}$ and n=5 $Ube2n^{MyeWT}$ tumors. Colocalization analysis was performed in CellSens using a manual threshold applied identically to all images and across both genotypes. Statistical significance was determined by unpaired two-tailed t-test with Welch's correction. *p < 0.05, **p < 0.01, ***p < 0.001.

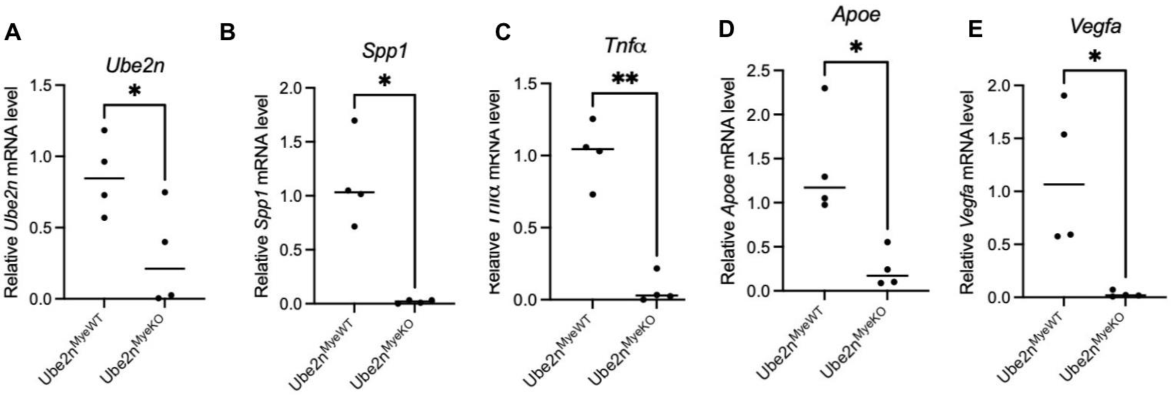


**Figure 3: UBE2N deletion in macrophages decreases Ube2n-dependent inflammatory and immunosuppressive genes.** (A-E) Relative mRNA expression of *Ube2n*, *Spp1*, *Tnfa*, *Apoe,* and *Vegfa* in either $Ube2n^{MyeKO}$ or $Ube2n^{MyeWT}$ BMDMs treated with 4-OHT for 72 hours then stimulated with TCM for 12 hours. Each point represents one independent bone marrow preparation. Technical replicates were averaged within each preparation before analysis. Data are shown as means with individual values. N= 4 independent bone marrow preparations per genotype. All data are normalized to the *Gapdh* and *Stx5a* housekeeping genes. Statistical significance was determined using an unpaired two-tailed t-test with Welch's correction. *p < 0.05, **p < 0.01, ***p < 0.001.

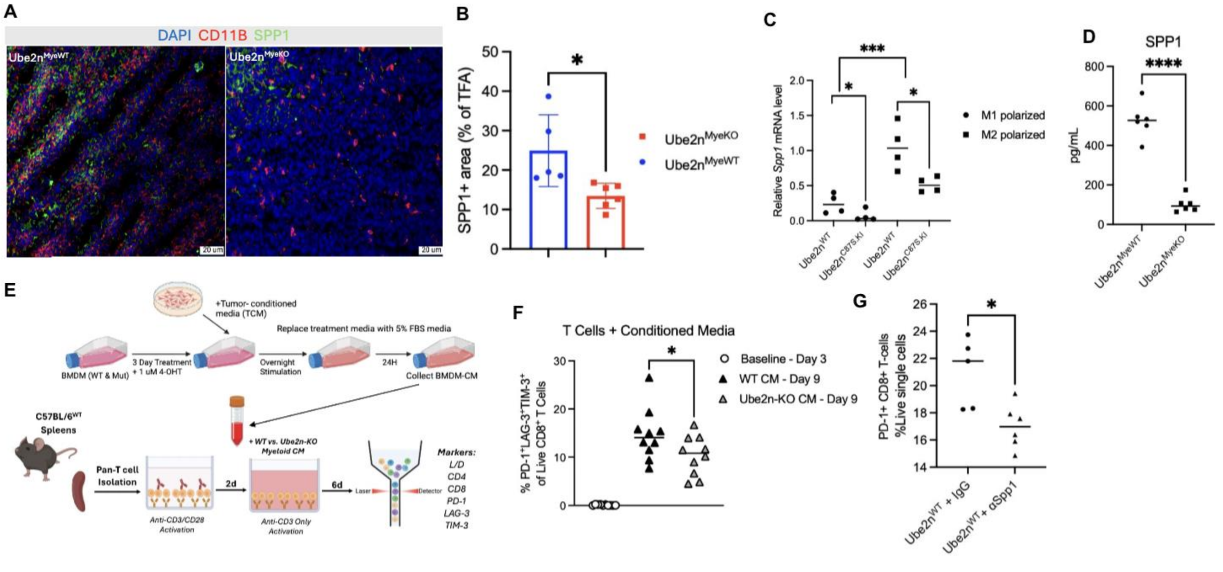

**Figure 4: Myeloid *Ube2n* supports SPP1 production, and myeloid-derived SPP1 alters CD8+ T cell checkpoint marker expression.** (A) Representative immunofluorescence images of tumor sections from $Ube2n^{MyeKO}$ and $Ube2n^{MyeWT}$ animals stained for CD11B (red), SPP1 (green), and DAPI nuclear counterstain (blue). Scale bar = 20 um. (B) Quantification of SPP1+ area normalized to TFA. Each point represents one tumor. Field-level measurements were averaged within each tumor. Comparisons were performed at the level of the animal. (C) Relative mRNA level of *Spp1* in M1- and M2-polarized $Rosa^{CreERT2}.Ube2n^{C87S.KI}$ and $Rosa^{CreERT2}.Ube2n^{WT}$ BMDMs treated with 4-OHT. Each point represents an independent culture. N=4 cultures per condition. Data are shown as means with individual values +/- SD. Statistical significance was determined using an unpaired two-tailed t-test with Welch's correction. (D) SPP1 protein concentration in BMDM culture media supernatant from 4-OHT treated $Ube2n^{MyeKO}$ and $Ube2n^{MyeWT}$ BMDMs. Each point represents an independent culture. N=6 cultures per genotype. (E) Scheme of T-cell:BMDM-CM experiment. (F) Frequency of PD-1+ LAG-3+ TIM-3+ cells among live CD8+ T cells at baseline (day 3) and after 6 days of culture (day 9) in either $Ube2n^{MyeKO}$ or $Ube2n^{MyeWT}$ BMDM-CM. Each point represents an individual culture well. (G) Frequency of PD-1+ CD8+ T cells among live single cells following co-culture with $Ube2n^{MyeWT}$ BMDMs in the presence of an IgG isotype control or SPP1-neutralizing antibody (aSPP1). Each point represents an individual culture well. Data are shown as individual values + SD. Statistical significance was determined using an unpaired two-tailed t-test with Welch's correction. *$p < 0.05$, **$p < 0.01$, ***$p < 0.001$, ****$p<0.0001$.
